# Plant–hummingbird pollination networks exhibit minimal rewiring after experimental removal of a locally abundant plant species

**DOI:** 10.1101/2022.10.05.511021

**Authors:** Kara G. Leimberger, Adam S. Hadley, Matthew G. Betts

**Affiliations:** Forest Biodiversity Research Network, Department of Forest Ecosystems and Society, Oregon State University, Corvallis, Oregon, USA; Biodiversity Section, Department of Natural Resources and Energy Development, Fredericton, New Brunswick, Canada

**Keywords:** partner switching, adaptive foraging, network dissimilarity, individual specialization, coextinction

## Abstract

1. Mutualistic relationships, such as those between plants and pollinators, may be vulnerable to the local extinctions predicted under global environmental change. However, network theory predicts that plant–pollinator networks can withstand species loss if pollinators switch to alternative floral resources (rewiring). Whether rewiring occurs following species loss in natural communities is poorly known because replicated species exclusions are difficult to implement at appropriate spatial scales.
2. We experimentally removed a hummingbird-pollinated plant, *Heliconia tortuosa*, from within tropical forest fragments to investigate how hummingbirds respond to temporary loss of an abundant resource. Under the *rewiring hypothesis*, we expected that niche expansions would decrease ecological specialization and reorganize the network structure (i.e., pairwise interactions).
3. We employed a replicated Before-After-Control-Impact experimental design and quantified plant–hummingbird interactions using two parallel sampling methods: observations of hummingbirds visiting focal plants (‘camera networks’, created from >19,000 observation hours) and pollen collected from individual hummingbirds (‘pollen networks’, created from >300 pollen samples). To assess hummingbird rewiring, we quantified ecological specialization at the individual, species, and network levels and calculated the amount of network-level interaction turnover (i.e., gain/loss of pairwise interactions). Leveraging our parallel network datasets, we also explored how sampling method influences apparent specialization.
4. *H. tortuosa* removal caused some reorganization of pairwise interactions but did not prompt large changes in specialization, despite the large magnitude of our manipulation (on average, >100 inflorescences removed in treatment areas of >1 ha). Although some individual hummingbirds sampled through time showed modest increases in niche breadth following *Heliconia* removal (relative to birds that did not experience resource loss), these changes were not reflected in species- and network-level specialization metrics. We also found that camera networks were more specialized than pollen networks, and that correlation between sampling methods was low.
5. Our results suggest that animals may not necessarily shift to alternative resources after losing an abundant food resource, even in species thought to be highly opportunistic foragers such as hummingbirds. Given that rewiring contributes to theoretical predictions of network stability, future studies should investigate why pollinators might not expand their diets after a local resource extinction.

## INTRODUCTION

Under global environmental change, a critical research challenge is to understand how anthropogenic stressors will alter species interactions (Burkle & Alarcón, 2011; Tylianakis *et al*., 2008; Walther, 2010). For instance, species may be lost from a community through numerous mechanisms as climate change intensifies (Dunn *et al*., 2009; Hegland *et al*., 2009). Climate change causes geographic range shifts to higher latitudes and elevations (Parmesan, 2006; Walther *et al*., 2002), but range shifts of mobile animals will likely outpace those of plants (Corlett & Westcott, 2013). Thus, pollinators undergoing range shifts may encounter communities lacking their original plant partners (Dunn *et al*., 2009). Further, climate change alters the timing of plant flowering, leading to temporal decoupling between pollinators and their preferred floral resources (Hegland *et al*., 2009; McKinney *et al*., 2012; Memmott *et al*., 2007). Over shorter time scales, extreme weather events – such as hurricanes – can also destroy floral resources that pollinators depend on (Temeles & Bishop, 2019). When species depend on each other, as in the mutualistic interactions between plants and pollinators, loss of one species may trigger the loss of additional species (Colwell *et al*., 2012; i.e., a coextinction cascade: Dunn *et al*., 2009).

However, much uncertainty still exists about whether species loss causes coextinction cascades within real-world mutualistic networks. Numerous simulation studies have investigated the extinct to which species removal triggers additional extinctions and have generally found that mutualistic networks should be able to withstand single species losses, except when a centrally connected partner is removed (Kaiser-Bunbury *et al*., 2010; e.g., Memmott *et al*., 2004; Traveset *et al*., 2017). However, replicated field experiments that test this prediction remain relatively rare, because species exclusions are logistically difficult to implement at appropriate spatial scales (Brosi *et al*., 2017; Brosi & Briggs, 2013; Kaiser-Bunbury *et al*., 2017; but see Lopezaraiza–Mikel *et al*., 2007; Timóteo *et al*., 2016).

One mechanism that may allow pollinators to cope with species loss is their ability to switch interaction partners (“rewiring” sensu Kaiser-Bunbury *et al*., 2010). This behavioural flexibility not only benefits the pollinator, but also sustains pollination services to co-occurring plants, theoretically preventing extinction cascades in mutualistic networks (e.g., Kaiser-Bunbury *et al*., 2010; Schleuning *et al*., 2016; Valdovinos *et al*., 2013; Vizentin-Bugoni *et al*., 2020). However, whether this behavioural mechanism can buffer natural communities from collapse remains poorly known. On the one hand, pollinators commonly switch plant partners in seasonal environments with a rotating menu of nectar sources (CaraDonna *et al*., 2017; e.g., Petanidou *et al*., 2008) and diet expansions have occurred in previous species removal experiments (e.g., Brosi & Briggs, 2013; Costa *et al*., 2018; Timóteo *et al*., 2016). On the other hand, flexibility may be limited due to constraints, such as morphological trait matching and/or interspecific competition (Brosi & Briggs, 2013; Vizentin-Bugoni *et al*., 2020). These constraints may be especially pronounced in mutualistic networks where pairwise interactions are primarily structured by traits, such as those between hummingbirds and flowering plants (Maruyama *et al*., 2014; Vizentin-Bugoni *et al*., 2014). Because high levels of rewiring may also be necessary to prevent network collapse after plant species loss (Schleuning *et al*., 2016), understanding any limits to behavioural flexibility is critical for realistic predictions about network robustness.

Here, we experimentally removed a flowering plant (*Heliconia tortuosa*, hereafter ‘*Heliconia’*) from tropical plant–hummingbird interaction networks to investigate the extent to which hummingbirds altered their foraging preferences – thus indicating a capacity for rapid interaction rewiring. Under this *‘rewiring hypothesis’*, hummingbirds add alternative resources to their diet to cope with losing a previously abundant floral resource. This niche breadth expansion should lead to reduced ecological specialization (hereafter ‘specialization’) and a reorganization of the underlying network structure (pairwise interactions). Alternatively, hummingbirds might not expand their diets, perhaps due morphological or behavioral constraints, yielding minimal changes in specialization and network structure.

To investigate this hypothesis, we examined specialization at multiple organizational scales (community, species, and individual) using two complementary approaches for quantifying networks: (i) a pollinator-centered dataset (pollen grains collected from individual captured hummingbirds, hereafter *‘pollen network’*) and (ii) a plant-centered dataset (observations of hummingbirds visiting plants, hereafter *‘camera network’*). To further our understanding of these parallel sampling methods, we also compared paired networks of each type (pollen *versus* camera, *N* = 20 pairs), expecting that pollen networks would be less specialized than pollen networks because they reflect the perspective of highly mobile pollinators moving across greater spatial scales (Jordano, 2016; Ramírez-Burbano *et al*., 2017). Studies that use multiple paired networks to compare sampling method are rare in the pollination literature (we located one study: Zhao *et al*., 2019) but essential for predicting how networks will respond to species loss (Bosch *et al*., 2009). Further, our study is one of relatively few replicated experiments testing how mutualistic networks respond to species loss (Biella *et al*., 2019; Brosi *et al*., 2017; Brosi & Briggs, 2013; Kaiser-Bunbury *et al*., 2017; <10 total studies: Lopezaraiza– Mikel *et al*., 2007; Timóteo *et al*., 2016) and the only one that combines relatively high replication, a large spatial scale of manipulation, and multiple network sampling approaches.

## MATERIALS AND METHODS

### Study system

We conducted this study within 14 forest fragments (‘sites’) surrounding the Las Cruces Biological Station in southern Costa Rica (Coto Brus Canton, Puntarenas Province, 8°47’
s7″ N, 82°57′32″ W). All fragments used in this study bordered cattle pastures and/or small plantations of shade-grown coffee and were likely created during a period of extensive forest clearing from 1960–1980 (Zahawi *et al*., 2015). More than 20 hummingbird and >150 flowering plant species have been documented in this study system, though not all species can be found in individual fragments (Borgella *et al*., 2001; Hadley *et al*., 2018; Morrison & Mendenhall, 2020). Elevation of sites ranged from ∼900–1500 m. Data were collected across three years (2016–2018) between February and May; these months reflect the transition period between dry and rainy seasons (Borgella *et al*., 2001; Zahawi *et al*., 2015).

### Experimental design

This experiment used a replicated Before-After-Control-Impact (BACI) design, which involves paired sites (here, ‘Control’ and ‘Treatment’) monitored across two experimental periods (here, ‘Pre’ and ‘Post’) (Fig. 1D). We removed *Heliconia* by temporarily covering the inflorescences with dark-colored bags (Fig. 1C). The number of inflorescences covered depending on fragment area, *Heliconia* density, and navigability of terrain; on average, we removed 172 ± 138 SD *Heliconia* inflorescences (median: 138, range: 8–526) within areas of 1.7 ± 1.9 SD ha (median: 1.25 ha, range: 0.27–7.49 ha). The covered inflorescences comprised 96% (± 5.5%) of the *Heliconia* resources encountered; two *Heliconia* plants remained uncovered to maintain plant species composition between experimental periods, and sometimes plants were inaccessible. In the control site, the disturbance of *Heliconia* removal was replaced with a survey of floral resources (except in the first year). Each experiment lasted nine days, with *Heliconia* removal starting at midday on Day 4. The following day (Day 5) was designated as a behavioral adjustment day during which no data were collected.

**Figure 1.**
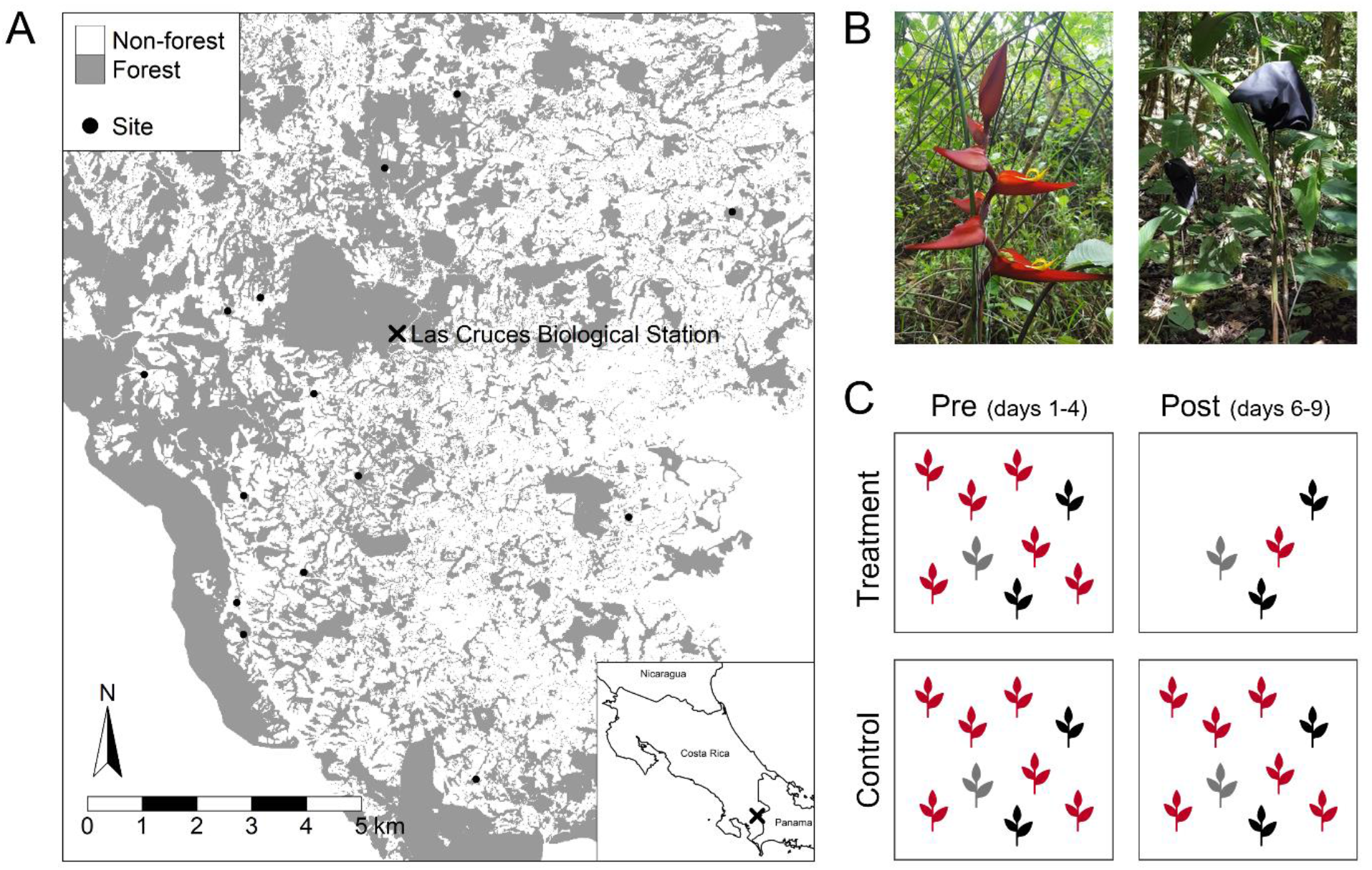
Within study sites (*N* = 14) surrounding the Las Cruces Biological Station in Costa Rica, we investigated the effects of removing a locally abundant flowering plant species (*Heliconia tortuosa*) using a Before-After-Control-Impact experimental design. **(A)** Study landscape, with forested areas shown in grey. The forest layer is a combination of areas digitized for a companion study and those digitized by Mendenhall & Wrona **(2018)**. Non-forested areas (white) are primarily pasture. **(B)** *Heliconia tortuosa* (left), the species of flowering plant that we temporarily removed by covering its inflorescences with dark bags (right). **(C)** The Before-After-Control-Impact design used in this experiment. Within a year (2016–2018), each site received either the *Heliconia* removal treatment (‘Impact’) or was assigned to be a Control. Treatments were reversed in alternate years for a total maximum sample size of 16 control replicates and 16 treatment replicates. Additionally, interactions between plants and hummingbirds were sampled during two experimental periods – Before (‘Pre’) and After (‘Post’). With this experimental design, the comparison of interest is the pre-to-post change in treatment replicates compared to the pre-to-post change in control replicates. Under the *rewiring hypothesis*, we expected that hummingbirds in treatment replicates would increasingly visit non-*Heliconia* plant species (grey and black icons) after removal of *Heliconia* (red icons). Note that two *Heliconia* plants per treatment site remained uncovered, which maintained plant species composition between experimental periods. *Photos provided by M. Donald*.

Due to limited site availability, we reused sites across the three study years. We randomly assigned initial treatments and statistically accounted for non-independence within sites with a random effect (see ‘*Statistical analysis’*, below). Sites used multiple times received the opposite treatment of the previous year, except when prevented by logistical considerations (especially challenging terrain: ∼20% of treatment assignments). Because most sites received multiple treatments, we refer to ‘treatment replicates’ and ‘control replicates’ (rather than ‘treatment sites’ and ‘control sites’). We repeated the *Heliconia* removal experiment 16 times, yielding a total of 16 control replicates and 16 treatment replicates. However, data from both experimental periods were not always available for each replicate, so some analyses had lower sample sizes (Table S1: *N*_max_: 15 control, 15 treatment; *N*_min_: 6 control, 5 treatment).

### Sampling plant–hummingbird interactions

We quantified plant–hummingbird interactions using two parallel sampling approaches. To construct pollen transport networks, we captured hummingbirds using mist nets (Days 1 and 9). Nets were opened 30 minutes after sunrise (to allow hummingbirds to forage on flowers before being captured) and remained opened for 2–5 hours. Capture effort and mist net locations remained constant across capture sessions (pre and post). As soon as captured hummingbirds were removed from the net, we sampled pollen grains from their beaks, nares, throats, and foreheads using glycerin-based fuchsin gel and mounted the sample onto a microscope slide (Kearns & Inouye, 1993; Maglianesi *et al*., 2015). Hummingbirds were then marked using aluminum tarsus bands, which enabled individual identification. If a bird was captured multiple times per capture session, we only included the first pollen sample in analysis (when collection time was available; *N* = 7 birds) and randomly chose one sample otherwise (*N* = 7 birds). We identified pollen grains using a light microscope and a pollen reference collection, which we created by opportunistically sampling flowers encountered during field work. Across all hummingbird pollen samples, we recorded 45 pollen morphotypes at varying degrees of taxonomic resolution. Thirty morphotypes were identified to species (*N* = 17), genus (*N* = 7), or family (*N* = 6); the remaining morphotypes were visually distinct but could not be confidently assigned to family. Of the 319 pollen samples collected from 292 individual hummingbirds, 307 samples (96%) had pollen and could be included in further analysis. All birds were handled in accordance with Oregon State University Animal Care and Use Protocols (ACUP #4655, #5015).

To construct camera networks, we used trail cameras (PlotWatcher Pro, Day 6 Outdoors) to directly observe hummingbirds visiting the flowers of 2–11 plant species per site (Days 2–4, 6–8). Many of these plants occurred naturally, but to each site we also added two floral arrays of potted plants (3–5 species/array). Floral arrays were established 3.7 ± 1.8 (mean ± SD) days before data collection to allow hummingbirds to discover the flowers (range: 0–8 days, median: 4 days). The arrays partially standardized plant species composition between paired sites and ensured that non-*Heliconia* flowers were available throughout the experimental period. On average, we obtained data from 10.3 ± 3.4 SD cameras per focal area (range: 2–17 cameras, median: 10.5 cameras). This effort was primarily concentrated at two ‘stations’ (>80% of cameras) that included the floral array of potted plants, one *Heliconia* plant whose inflorescence remained uncovered, and any naturally occurring flowering plants within a ∼5 m radius. Stations were located 60 ± 44 m apart (mean ± SD). All plant species observed were known or suspected to be visited by hummingbirds, and nearly all were in the forest understory (two overstory species, genus *Erythrina*, were sampled when accessible). Trail cameras took a photograph every second and combined these images into a time-lapse video. We then reviewed videos using machine-learning software (MotionMeerkat: Weinstein, 2015) or by watching the video at 3–4 times normal speed (GameFinder software, Day 6 Outdoors). The overall dataset comprised 20,735 video hours (Table S2). Analyses related to the experiment used 19,870 video hours (6475 hummingbird visits), while analyses of sampling method used 9,603 video hours (videos from the ‘pre’ period only).

### Constructing interaction networks

We constructed quantitative pollen transport networks by using the number of samples with a given pollen morphotype as the interaction frequency (Maglianesi *et al*., 2015; Ramírez-Burbano *et al*., 2017). For camera networks, we used the visitation rate per plant species as the interaction strength (i.e., number of hummingbird visits/number of video hours). We did not exclude visits in which hummingbirds obtained nectar by bypassing a flower’s reproductive structures (i.e., nectar robbing: Irwin *et al*., 2010) because we were interested in how *Heliconia* removal altered hummingbird foraging behavior, regardless of the pollination outcome. Because one network metric (*d*’, see ‘*Specialization’* below) can only be calculated for integers, we multiplied all interaction frequencies by 10,000 and rounded to the nearest whole number (Ballantyne *et al*., 2015; e.g., Pauw & Stanway, 2015). When estimating sampling completeness (see *‘Interaction turnover’* below), we used the raw number of hummingbird visits rather than hourly visitation rate.

### Response variables

#### Specialization

Under the rewiring hypothesis, we expected hummingbirds to become less ecologically specialized with local extinction of *Heliconia*. To quantify specialization at the individual level, we used the number of pollen morphotypes per individual hummingbird (Maglianesi *et al*., 2015; Timóteo *et al*., 2016). We analyzed individual specialization for all hummingbirds captured (‘all individuals’) and for individual hummingbirds caught during both capture sessions within a replicate (‘recaptures only’, *N* = 27 individuals).

At the species and network level, we calculated ecological specialization as *H*_2_’ (two-dimensional Shannon diversity), which indicates the degree of reciprocal specialization between interaction partners (Blüthgen *et al*., 2006). We also calculated two species-level metrics from the hummingbird perspective: standardized Kullback-Leibler distance (*d*’: Blüthgen *et al*., 2006) and the Species Specificity Index (SSI: Julliard *et al*., 2006). Conceptually, *d’* describes the exclusiveness of interactions (niche partitioning), though it may underestimate specialization if a rare and truly specialized species relies on a commonly visited partner (Blüthgen, 2010). The Species Specificity Index reflects niche breadth and is the coefficient of variation, calculated from visitation rates to each plant species (Julliard *et al*., 2006). Compared to other readily available network metrics, these three quantitative indices are robust to changes in sampling intensity (i.e., observations per species) and network size (Fründ *et al*., 2016). All indices range from 0 to 1, with higher values indicating higher specialization (i.e., high niche exclusiveness and/or low niche breadth). We averaged values of *d’* and SSI across (i) all hummingbird species and (ii) across two hummingbird species expected to rely most strongly on *H. tortuosa*, green hermits (*Phaethornis guy*) and violet sabrewings (*Campylopterus hemileucurus*) (Borgella *et al*., 2001; Taylor & White, 2007). Network metrics were calculated using the ‘networklevel’ and ‘specieslevel’ functions within the ‘bipartite’ R package (Dormann, 2011; Dormann *et al*., 2009).

#### Interaction turnover

Because specialization indices do not directly reflect changes in the identity of pairwise interactions, we also quantified interaction turnover between paired pre and post networks (Fig. 4.1). Interaction turnover describes the extent to which individual links are gained or lost over time and can be quantified using Whittaker’s beta diversity index (also called Sørensen dissimilarity) following Poisot *et al*. (2012) and Fründ (2021):

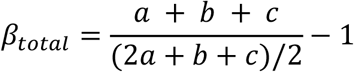

where *a* is the number of interactions shared between networks, *b* is the number of interactions present only in the ‘pre’ network, and *c* is the number of interactions present only in the ‘post’ network. To help identify the mechanisms driving any potential differences in interaction turnover, we also partitioned *β*_*total*_ into its additive subcomponents: (i) interaction turnover due to species gain or loss (‘species turnover’) and (ii) interaction turnover due to interaction reorganization among the species shared by both networks (Poisot *et al*., 2012; see also Fig. S1). Species turnover includes situations where a hummingbird species was only observed during one period, or a plant species received visits in one period but not the other. Therefore, if *Heliconia* removal causes hummingbirds to forage on completely novel resources (i.e., plants previously unused by any other hummingbird species), this change would be encapsulated by the species turnover component. The rewiring component reflects a more subtle reshuffling of interactions among hummingbird species, especially among common species more likely to be detected in both networks over time. Here we focus on binary indices of interaction turnover, which allow a more straightforward interpretation of the partitioned components than frequency-based indices (Fründ, 2021), though results were qualitatively similar with a quantitative metric (Bray-Curtis dissimilarity). We calculated interaction turnover and its components using the ‘betalinkr’ function within the ‘bipartite’ R package (Fründ, 2021). Indices range from 0 to 1, with higher values indicating higher turnover.

Because differences in network size and sampling effort can influence estimates of interaction turnover (Poisot *et al*., 2012), we also estimated the sampling completeness of each network. First, we estimated interaction richness as the bias-corrected Chao 1 index, calculated using the ‘estimateR’ function of the ‘vegan’ R package (Oksanen *et al*., 2020). Applying this approach to interaction networks is analogous to estimating species richness, except that it uses the vectorized interaction matrix; each pairwise interaction becomes a row (‘species’) with an associated interaction frequency (‘abundance’) (Jordano, 2016). We divided observed interaction richness by the estimated interaction richness to calculate sampling completeness (CaraDonna *et al*., 2017; Ramírez-Burbano *et al*., 2017; Robinson & Strauss, 2020).

### Statistical analysis

We conducted all statistical analyses in R version 4.2.0 (R Core Team, 2022). We modeled network indices and sampling completeness using beta regression after transforming the response variable to exclude 0 and 1 (as recommended by Cribari-Neto & Zeileis, 2010). For individual-level specialization, measured as the number of pollen morphotypes per hummingbird, we used a truncated generalized Poisson distribution. This distribution accommodated the lack of zeroes in the dataset and yielded lower AICc values than models with either the Poisson or negative binomial families. We implemented all models in the ‘glmmTMB’ package (Brooks *et al*., 2017) and verified that model assumptions were met by visually inspecting plots produced by ‘DHARMa’ (Hartig, 2020). We also checked for highly influential replicates using Cook’s distance and DFBETAS in the ‘influence_mixed’ package. To interpret results, we calculated treatment contrasts and 95% confidence intervals using ‘emmeans’ (Lenth, 2020).

#### Responses to Heliconia removal

When modelling response variables measured during both experimental periods, we included a statistical interaction between *Heliconia* removal (control *versus* treatment) and experimental period (pre *versus* post), because we expected that pre-to-post change would be greater in treatment replicates than control replicates. Analyses of network dissimilarity, calculated from paired pre-to-post networks, only included a predictor variable for *Heliconia* removal. We accounted for re-use of the same site across multiple years with a random intercept for Site. When applicable, we included a random intercept for Replicate, meant to pair ‘pre’ and ‘post’ observations within a replicate. When modeling individual-level specialization of recaptured hummingbirds, we also included a random intercept for Bird to pair ‘pre’ and ‘post’ observations within a bird (and account for potential non-independence of foraging behavior within individuals). Sample sizes varied with sampling method and response variable, being highest for camera networks and lowest for individual recaptures (Table S1). Analyses of pollen networks had intermediate sample sizes because low hummingbird capture numbers sometimes produced single-species networks or networks with unconnected compartments; these networks were excluded from analysis.

#### Comparison of sampling methods

We only used data from the ‘pre’ period when analyzing network sampling methods, thus removing any influence from our removal experiment. First, we used beta regression to statistically investigate how sampling method influenced our three metrics of ecological specialization (*H*2’, *d*’, and SSI). Each model included a predictor variable for network type (pollen *versus* camera) and a random effect for Replicate, which paired networks sampled using different methods (*N* = 20 network pairs). Second, we used beta regression to examine the strength of the relationship between paired pollen and camera networks. Finally, to gain insight into overall pattern of plant–hummingbird interactions in this study system, we combined observations across sites and years into one network per sampling method (pollen *versus* camera meta-networks, Fig. 1).

## RESULTS

### Normal visitation patterns

Combining plant–hummingbird interactions across sites and years, we observed 16 hummingbird species and 39 pollen morphotypes in the pollen meta-network (Fig. 2A) and 17 hummingbird species and 32 plant species in the visitation meta-network (Fig. 2B). Green hermits and violet sabrewings accounted for 58% of the pollen samples containing *Heliconia* (103 of 177 samples) and 84% of *Heliconia* visits (889 of 1055 visits). Of the pollen samples collected from green hermits and violet sabrewings, 99% included *Heliconia* (103 of 104 samples). Fifty-four percent of these pollen loads with *Heliconia* were either monospecific (18 of 104 samples; 17%) or contained only one additional pollen type (38 of 104 samples; 37%).

**Figure 2.**
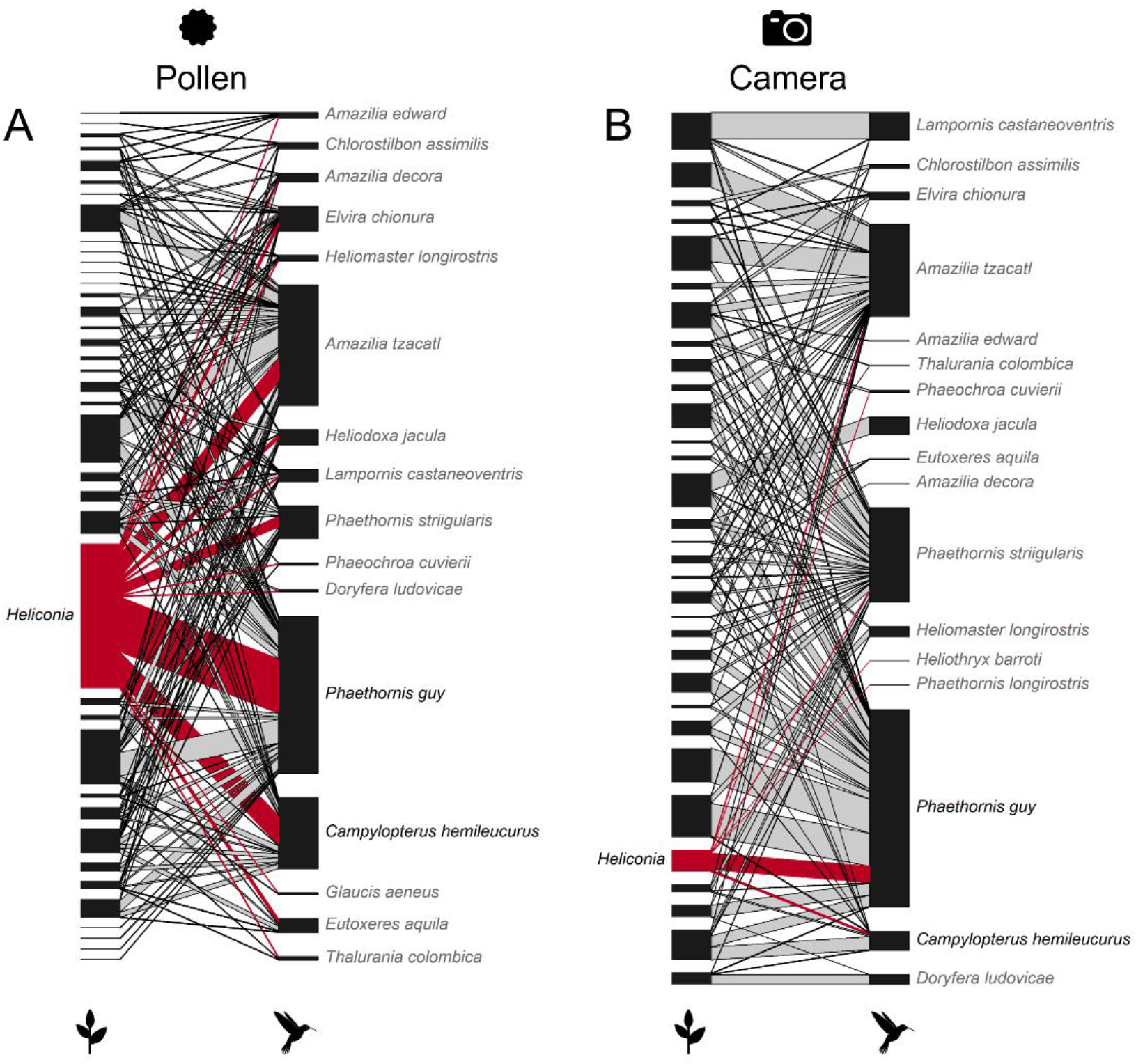
Meta-networks of plant–hummingbird interactions created by combining observations across three field seasons (2016-2018) and 14 sites surrounding the Las Cruces Biological Station in southern Costa Rica. The pollen network **(A)** was created by identifying pollen grains sampled from captured hummingbirds (228 pollen samples), and the camera network **(B)** was created from trail cameras observing hummingbirds visiting focal plants (15,119 video hours). Connections to *Heliconia tortuosa* are highlighted in red. For pollen networks, the interaction frequency is the number of pollen samples containing a given pollen morphotype. For camera networks, the interaction frequency is the number of hummingbird visits per hour of video observation. The height of each rectangle represents the overall observation frequency for that species (larger rectangle = more observations), while the width of the connecting links reflects relative interaction frequency between partners (wider link = higher interaction frequency). Note that two hummingbird species, the green hermit (*Phaethornis guy*) and violet sabrewing (*Camplyopterus hemileucurus*), are linked most strongly to *Heliconia*.

**Figure 3.**
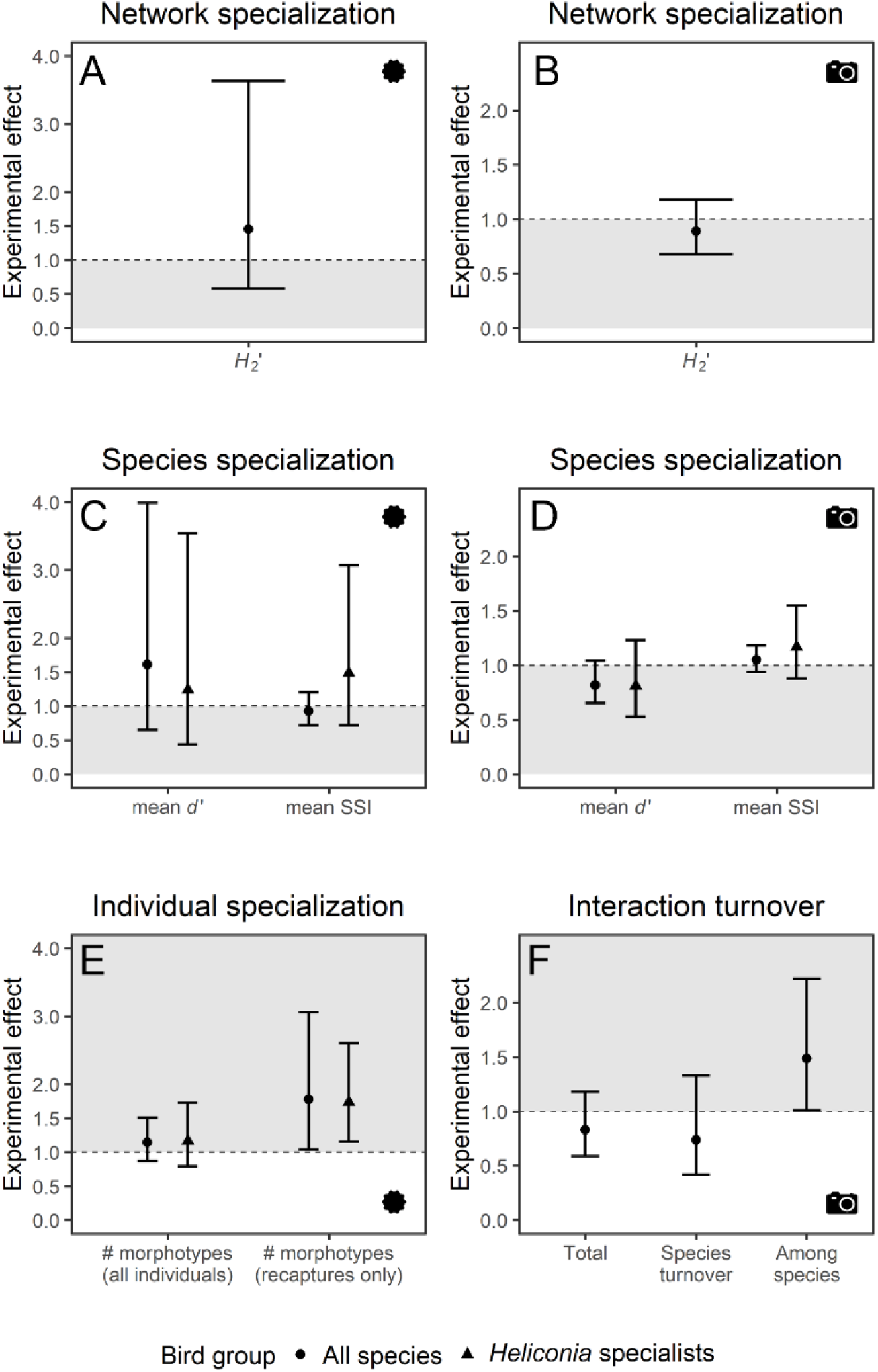
Effects of experimental *Heliconia* removal on plant–hummingbird interactions measured with two sampling methods: pollen grains from captured hummingbirds (left column) and video observations of hummingbirds visiting plants (right column). Following the Before-After-Control-Impact experimental design, the *y*-axis reflects the pre-to-post change in treatment replicates relative to pre-to-post change in control replicates (A-E), or treatment replicates relative to control replicates (F). Estimates and 95% confidence intervals were calculated using the ‘contrasts’ function of ‘emmeans’ (Lenth, 2020). No change is indicated by the dashed line at 1, and grey shading indicates support of the *rewiring hypothesis* (i.e., regions of decreased specialization or increased turnover). **(A-B)** Network*-*level specialization, quantified as *H*_2_’ (two-dimensional Shannon diversity, indicating niche overlap). **(C-D)** Species-level specialization, quantified as *d*’ (standardized Kullback-Leibler distance, indicating niche overlap) and Species Specificity Index (coefficient of variation, indicating niche breadth). *Heliconia* specialists include two hummingbird species: green hermits (*Phaethornis guy*) and violet sabrewings (*Campylopterus hemileucurus*). **(E)** Individual-level specialization, quantified as the number of pollen morphotypes per hummingbird, both for all individuals and individuals captured during both experimental periods (pre and post). **(F)** Total interaction turnover between pre and post networks and its additive subcomponents: turnover due to species gain and/or loss, and turnover due to interaction reshuffling among species shared by both networks. Results are shown for Whitaker’s beta diversity, a binary metric of network dissimilarity.

Across all hummingbird species, individual birds carried 2.6 ± 1.4 (mean ± SD) pollen morphotypes at the time of capture (range: 1-8, median: 2).

### Responses to Heliconia removal

#### Specialization

Under the *rewiring hypothesis*, we expected hummingbirds to expand their niche breadths after *Heliconia* removal. However, this hypothesis only received support when examining the hummingbirds captured during both experimental periods; on average, the pre-to-post change in pollen morphotypes per recaptured hummingbird was 1.78 times greater in treatment replicates than control replicates (95% CI: 1.04–3.06 times, Fig. 2E). A similar pattern emerged when analyzing recaptured *Heliconia* specialists (Fig. 2E, Table S3). When including all individuals captured, we did not find evidence of an experimental effect (treatment *x* period interaction, all species: *z* = 0.99, *P* = 0.32, Fig. 2E, Table S4). Similarly, we did not find that *Heliconia* removal led to statistically significant differences in any network-level (*H*_2_’) or species-level (*d*’, SSI) metrics of ecological specialization, even when examining species-level metrics averaged across *Heliconia* specialists (Fig. 2A-D, Tables S5-S6).

#### Interaction turnover

Because sampling differences can influence estimates of interaction turnover, we estimated the pre-to-post sampling completeness for each network. For camera networks, estimated sampling completeness was high (mean ± SD: 91 ± 11%, Table S7) and pre-to-post changes in sampling completeness were similar between control and treatment replicates (Table S8). However, consistent changes were not observed in pollen networks, which were also sampled less thoroughly (mean ± SD: 50 ± 19%, Table S7). Therefore, we only analyzed interaction turnover for camera networks, after also removing two outlier replicates (Fig. S2).

When examining overall interaction turnover, we did not detect pre-to-post differences in the networks from control *versus* treatment replicates (*z* = −1.08, *P* = 0.28, Fig. 2F, Table S9). The species turnover component, reflecting changes in species composition, also did not show an effect of *Heliconia* removal (*z* = −1.05, *P* = 0.29, Fig. 2F, Table S9). However, the rewiring subcomponent — representing interaction reorganization among species present during both experimental periods — was, on average, 1.49 times greater in treatment replicates (95% 1.01–2.22 times, Fig. 2F, Table S10).

### Comparison of sampling methods

On average, camera networks reflected 3.05 times more network-level specialization than pollen transport networks (95% CI: 2.08–4.48 times, Fig. 4A, Tables S11-S12). Results were qualitatively similar for the species-level specialization metrics (*d*’ and SSI) calculated from the hummingbird perspective (Fig. 4C, E, Tables S11-12). Pollen networks were weakly correlated with camera networks sampled during the same time period at the same site (e.g., *H*_2_’: *z* = 0.46, *P* = 0.64, pseudo-*R*^2^ = 0.014, Fig. 2B, Table S13).

**Figure 4.**
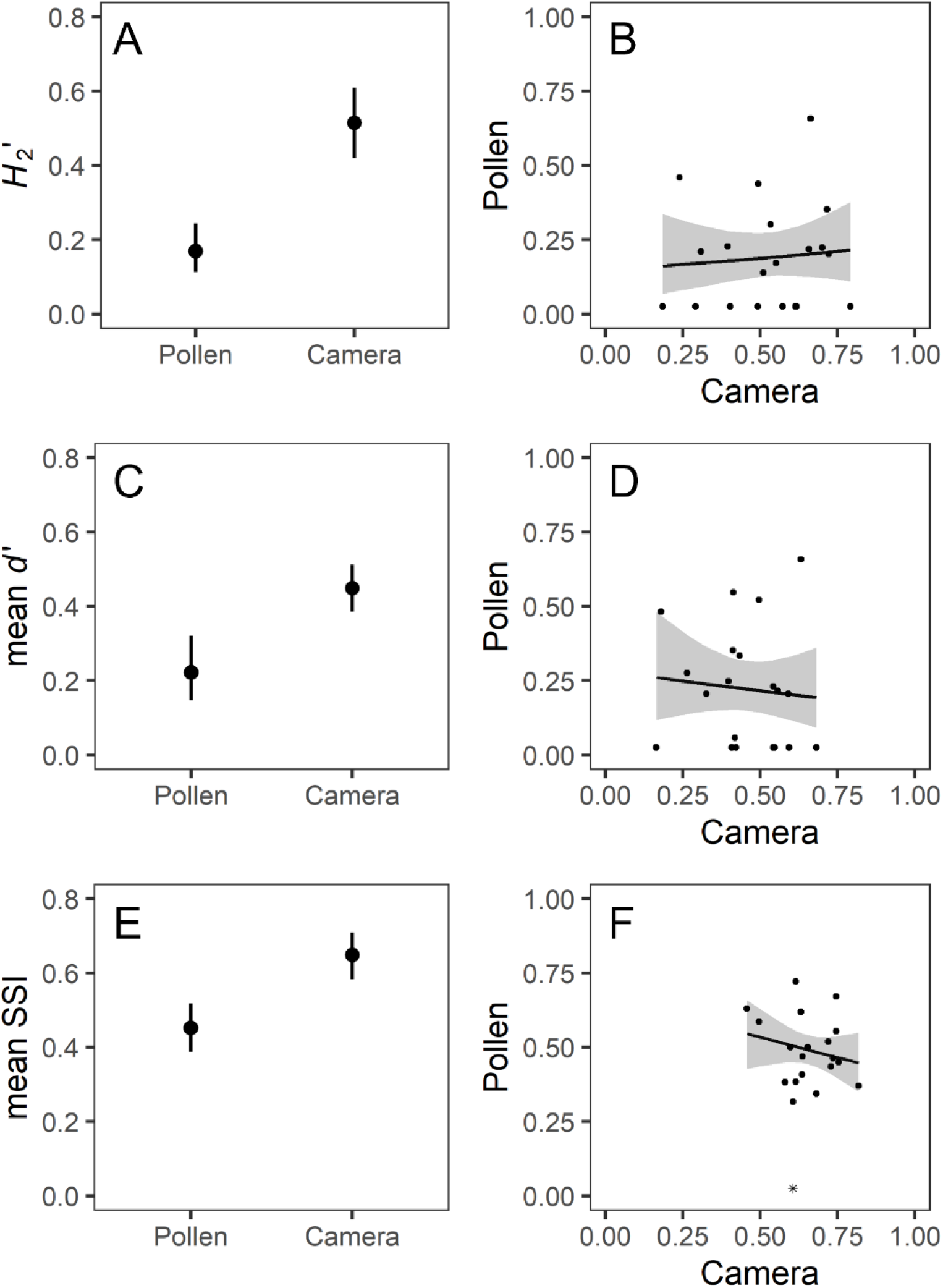
Comparisons of the two methods used to sample plant–hummingbird interactions in this study: pollen samples collected from captured hummingbirds (‘Pollen’) and direct observations of hummingbirds visiting focal plants (‘Camera’). **Left column:** Estimated effects of sampling method on three metrics of ecological specialization (*H*_2_’, *d*’, and Species Specificity Index), analyzed using the networks from each site and year. Estimated means are presented alongside 95% confidence intervals. Species-level metrics (*d*’, SSI) reflect the hummingbird perspective, averaged across all species. **Right column:** The estimated relationship between pollen and camera networks from the same site and year, calculated from beta regression models, alongside 95% confidence bands and raw data points. The analysis of SSI excludes one outlying data point (asterisk symbol in F).

## DISCUSSION

We experimentally simulated the local extinction of a hummingbird-pollinated flowering plant species to evaluate the extent to which interactions rapidly reorganize following species loss. Under the *rewiring hypothesis*, we expected that hummingbirds would increase their foraging on alternative resources, leading to niche expansions and increased niche overlap among species. Although we found some evidence for interaction restructuring, these changes appeared insufficiently large to alter most metrics of ecological specialization, lending limited support to the *rewiring hypothesis*.

When examining individual specialization, we only found evidence for an experimental effect for hummingbirds captured during both experimental periods, representing 9% of all individuals sampled for pollen. However, the effect did not arise from a niche expansion in treatment replicates; although hummingbirds carried an estimated 2.2 pollen morphotypes (95% CI: 1.7–2.9) before *Heliconia* removal and 2.8 pollen morphotypes after *Heliconia* removal (95% CI: 2.2–3.7), this increase was not statistically significant. Rather, the modest increase in treatment replicates was coupled with an *opposite* pattern in the control replicates (i.e., fewer pollen types over time, Fig. S3). Perhaps recaptured hummingbirds remained in focal areas throughout the experiment, so treatment birds were forced to maintain (or expand) their niche breadth to cope with *Heliconia* loss, relative to control birds that were simply responding to phenological changes (e.g., more *Heliconia* plants coming into bloom over time, and less reliance on other plant species).

Pollen networks analyzed at the species and network level also showed no evidence of reduced specialization after loss of a locally abundant interaction partner. This finding was somewhat unexpected, given the magnitude of our manipulation – both in terms of area covered (1.7 ± 1.9 ha) and average resources removed (96 ± 5.5% of *Heliconia* inflorescences). Further, *Heliconia* appeared to be a core diet component for its primary visitors, green hermits and violet sabrewings, nearly all of which were carrying *Heliconia* pollen grains when captured (see also Borgella *et al*., 2001). The lack of large-scale network changes also runs counter to several published examples of experimental species removals in mutualistic networks. In these studies, pollinators generally exhibited behavioral changes that resulted in increased niche overlap among species, as quantified through network-level specialization metrics (Brosi *et al*., 2017; Goldstein & Zych, 2016; Kaiser-Bunbury *et al*., 2017). However, *Heliconia* removal did appear to elicit changes in pairwise interactions within camera networks. This effect did not result from interactions gained or lost due to species turnover, but rather from interaction reorganization among plant and hummingbird species present in both experimental periods. We suspect that these subtle changes may have been due to fluctuations in the strength of interspecific (or intraspecific) competition, which can lead to rearrangement of foraging patterns (Brosi & Briggs, 2013).

We suggest this interaction reorganization did not emerge in the specialization indices because the magnitude of rewiring was relatively small, and network metrics can be insensitive to shifts in underlying interactions. For instance, after the dominant ant species was removed from a seed dispersal network, interactions were redistributed among the remaining species and not reflected by network-level specialization metrics (Timóteo *et al*., 2016). In the context of our experiment, there are several potential reasons why we observed minimal rewiring. One possibility is that plant–hummingbird interactions were constrained in some way, perhaps due to trait matching between hummingbird bills and flower corollas (Maruyama *et al*., 2014; Vizentin-Bugoni *et al*., 2014). Dominance relationships among (and within) hummingbird species could also limit the extent of interaction reshuffling (Carpenter *et al*., 1993; Stiles & Wolf, 1970; Temeles *et al*., 2005). Because dissimilarity metrics do not provide information about *which* interactions changed (Fründ, 2021), we cannot say whether some hummingbird species rewired more than others or if certain plant species were preferentially visited after resource removal (e.g., Biella *et al*., 2019). Further analysis could explore this possibility by incorporating characteristics known to structure plant–hummingbird interactions, such as hummingbird foraging strategy, floral corolla shape, and/or floral caloric value (see Leimberger *et al*., 2022 for review). Additionally, rewiring may not be the only mechanism that hummingbirds use to cope with resource loss. For one, highly mobile vertebrate pollinators can presumably still access preferred resources after an extirpation within part of their home range. If hummingbirds were able to access unmanipulated *Heliconia* plants with relative ease, extreme niche shifts may have been unnecessary over the short time scale of our experiment. Additionally, hummingbirds have physiological adaptations, such as torpor, that might have temporarily reduced their reliance on floral resources.

Comprehensively sampling plant–pollinator interactions at the community level is challenging (Jordano, 2016), and we acknowledge that our methodological approach has some limitations. For pollen networks, we cannot exclude the possibility that hummingbirds were transporting pollen from broad geographic extents, therefore introducing unwanted variability (Jordano, 2016; Ramírez-Burbano *et al*., 2017; Souza *et al*., 2021). Additionally, pollen networks relied on capturing enough hummingbirds (and sampling enough interactions) to create meaningful networks; low capture numbers in some sites limited our overall sample size for pollen network analysis, and estimated sampling completeness for pollen networks was low and highly variable (mean ± SD: 50 ± 19%). Nevertheless, we expect the limitations of pollen-based networks to be somewhat counterbalanced by their enhanced capacity to represent the hummingbird perspective. This advantage could be especially important within a multi-layered forest environment that includes many flowering lianas and epiphytes (Rocca & Sazima, 2013; e.g., passionflowers, bromeliads: Snow, 1982). Indeed, canopy floral resources were rarely represented by our camera arrays due to the additional resources needed to climb trees in a tropical forest. However, our camera networks do represent an intensive observational effort within the forest understory (generally 12 hours/day per focal plant, totaling >19,000 hours), leading to high estimates of sampling completeness for the plants sampled (>90%, on average). We also emphasize that – despite the various limitations inherent to network sampling – our BACI experimental design focuses on the *changes* between the pre and post experimental periods. Thus, even if we did not sample all interactions, we still are able to quantify changes in interactions among the plants we did sample.

Our networks also allowed us to compare two parallel methods for sampling plant– pollinator interactions: pollen loads sampled from captured hummingbirds *versus* direct observations of hummingbirds visiting focal plants. We found higher specialization in the camera networks, consistent with previous comparisons of these network sampling approaches (avian pollinators: Ramírez-Burbano *et al*., 2017; Zanata *et al*., 2017; insect pollinators: Bosch *et al*., 2009; but see Zhao *et al*., 2019; Souza *et al*., 2021). In our study, differences between sampling methods could arise because camera networks included visits where hummingbirds bypassed the plant’s anthers (robbing was observed in 11% of hummingbird visits). However, excluding nectar robbers from camera networks is expected to *increase* network-level specialization (Maruyama *et al*., 2015) – leading to more extreme differences than we report here. Another potential explanation for the discrepancy between methods is that the pollinator-centric approach sampled a larger spatial extent than focal plant observations (Jordano, 2016; Ramírez-Burbano *et al*., 2017). Additionally, our pollen networks have a coarser taxonomic resolution than our camera networks, as pollen grains were identified to morphotype based on visual characteristics. Therefore, multiple plant species might be represented by single pollen morphotypes, artificially increasing niche overlap among bird species. Intriguingly, Zhao *et al*. (2019) fine-tuned their pollen identifications by incorporating elevational and phenological information, finding that plant-insect camera networks were *less* specialized than their pollen network counterparts. To our knowledge, all existing comparisons of visitation and pollen networks have visually identified pollen based on morphological features, but more advanced techniques (i.e., DNA metabarcoding: Bell *et al*., 2017) would strengthen future investigations.

## CONCLUSIONS

Overall, our results suggest that the stabilizing effects of pollinator foraging flexibility might be relatively modest in real-world mutualistic networks, possibly operating alongside other mechanisms that enable short-term species persistence after sudden resource loss (e.g., high mobility or energy-saving adaptations). We did not investigate whether hummingbirds might have incorporated alternative resources over longer time periods (>4 days). However, if rewiring remains limited over extended time scales, then local extinctions could potentially trigger network collapse. Long-term studies of plant–pollinator interactions at range boundaries, across elevational gradients, and/or after extreme weather events cause sudden resource loss could help us better understand the behavioral flexibility of pollinators in the face of global environmental change.

## AUTHOR CONTRIBUTIONS

ASH and MGB conceived the ideas and designed methodology; KGL, ASH, and MGB collected the data; KGL analysed the data; KGL led the writing of the manuscript. All authors contributed critically to the drafts and gave final approval for publication.

## DATA AVAILABILITY STATEMENT

Data will be made available through the Dryad Digital Repository. Code for all analyses will be archived through Zenodo and is currently available at: github.com/kleimberger/Heliconia_removal_Network_rewiring

## ACKNOWLEDGEMENTS

This material is based upon work supported by the National Science Foundation (NSF) Graduate Research Fellowship Program under Grant No. 1840998 to KGL. Funding for field work was provided by NSF-DEB-1457837 to MGB and ASH, with additional support to KGL from the American Ornithological Society and Sigma Xi. We are also grateful to the Las Cruces Biological Station and the private landowners who allowed us to work on their property. Field assistance was provided by Jorge Araya Paniagua, Michael Atencio Picado, Marion Donald, Dustin Gannon, Jessica Greer, Matt Hadley, Urs Kormann, Daniel Moreno, Mauricio Paniagua Castro, Ignacio Ruiz Ilama, Esteban Sandi Paniagua, Andrés Solorzano Vargas, Laura Sutcliffe, and Felipe Torres Vanegas. Videos from trail cameras were reviewed with the help of Marion Donald, Colleen Franklin, Jessica Greer, Zoe Griffith, Elena Hart, Billy Hilgert, Zachary Kendall, Jeremy Lee, Ana Medina Roman, Briley Mullin, Amber Newell, Thanh Nguyen, Zoe Sallada, Allison Simoni, Andrew Stokes, Claire Woods, and Mel Xiao. The pollen reference collection was developed with the help of Jessica Greer, Amber Newell, and Andy Moldenke. Amber Newell provided the pollen identifications for this study. Mark Novak and Berry Brosi provided feedback on an earlier version of the manuscript. Icons within figures are from The Noun Project (creators Zach Bogart [pollen grain], Stanislav Levin [camera], alerma [plant] and Agne Alesiute [hummingbird]).

