## Supplemental Material for "Plant–hummingbird pollination networks exhibit minimal rewiring after experimental removal of a locally abundant plant species"

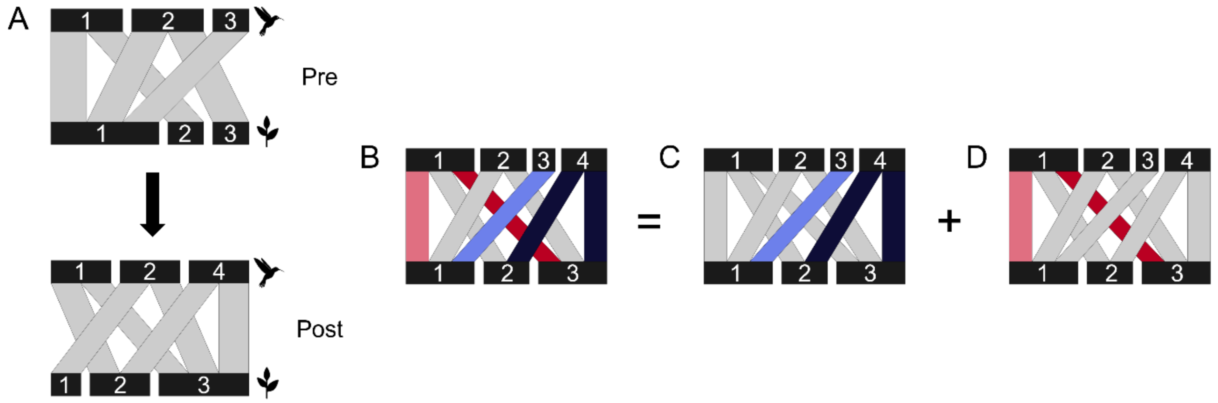


**Figure S1.** Visualization of interaction turnover in an example network with three hummingbird species and three plant species. **(A)** From the 'pre' period to the 'post' period, one hummingbird species (No. 3) is lost, and another hummingbird species (No. 4) is gained. Hummingbird species No. 1 switches partners, from plant species No. 1 to plant species No. 3. **(B-D)** Networks in A, combined to illustrate total interaction turnover (B) and its additive subcomponents: interaction turnover due to species gain or loss (C, blue links) and interaction turnover among species shared by both networks (D, red links). Darker links represent interaction gain; lighter links represent interaction loss.

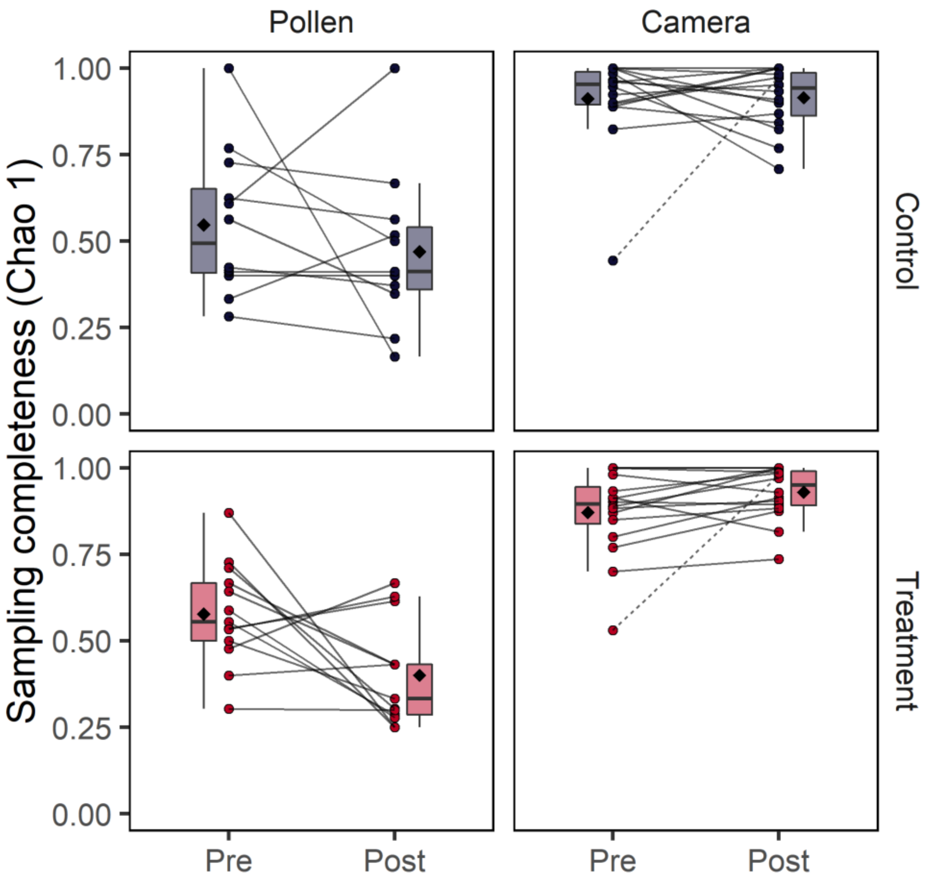


**Figure S2.** Estimated sampling completeness for each network, split across the two sampling methods used in this study: pollen samples from captured hummingbirds ('pollen') and direct oboservations of hummingbirds visiting flowers ('camera). Prior to analyzing interaction turnover for camera networks, we removed the two replicates with exceptionally large pre-to-post differences in sampling completeness (right column, dotted lines).

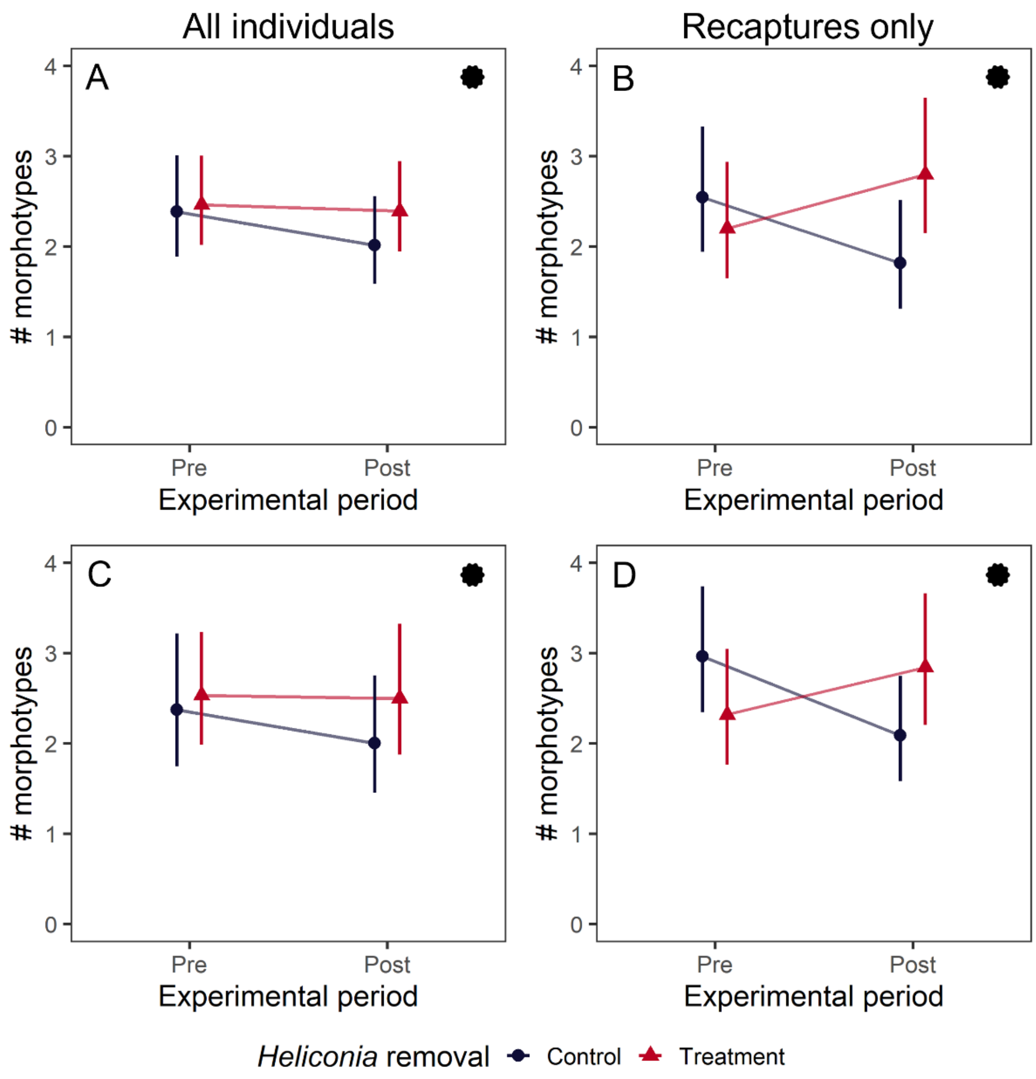


**Figure S3.** Effects of experimental *Heliconia* removal on individual-level specialization, measured as the number of pollen morphotypes sampled from captured hummingbirds. Results are shown for all hummingbirds captured (left column) and the subset of individuals captured during both experimental periods (right column). Estimated marginal means from GLMMs are presented alongside 95% confidence intervals. Conceptually, a treatment effect is indicated by non-parallel lines. (A-B) Results for all hummingbird species (C-D) Results for *Heliconia* specialists, green hermits (*P. guy*) and violet sabrewings (*C. hemileucurus*).

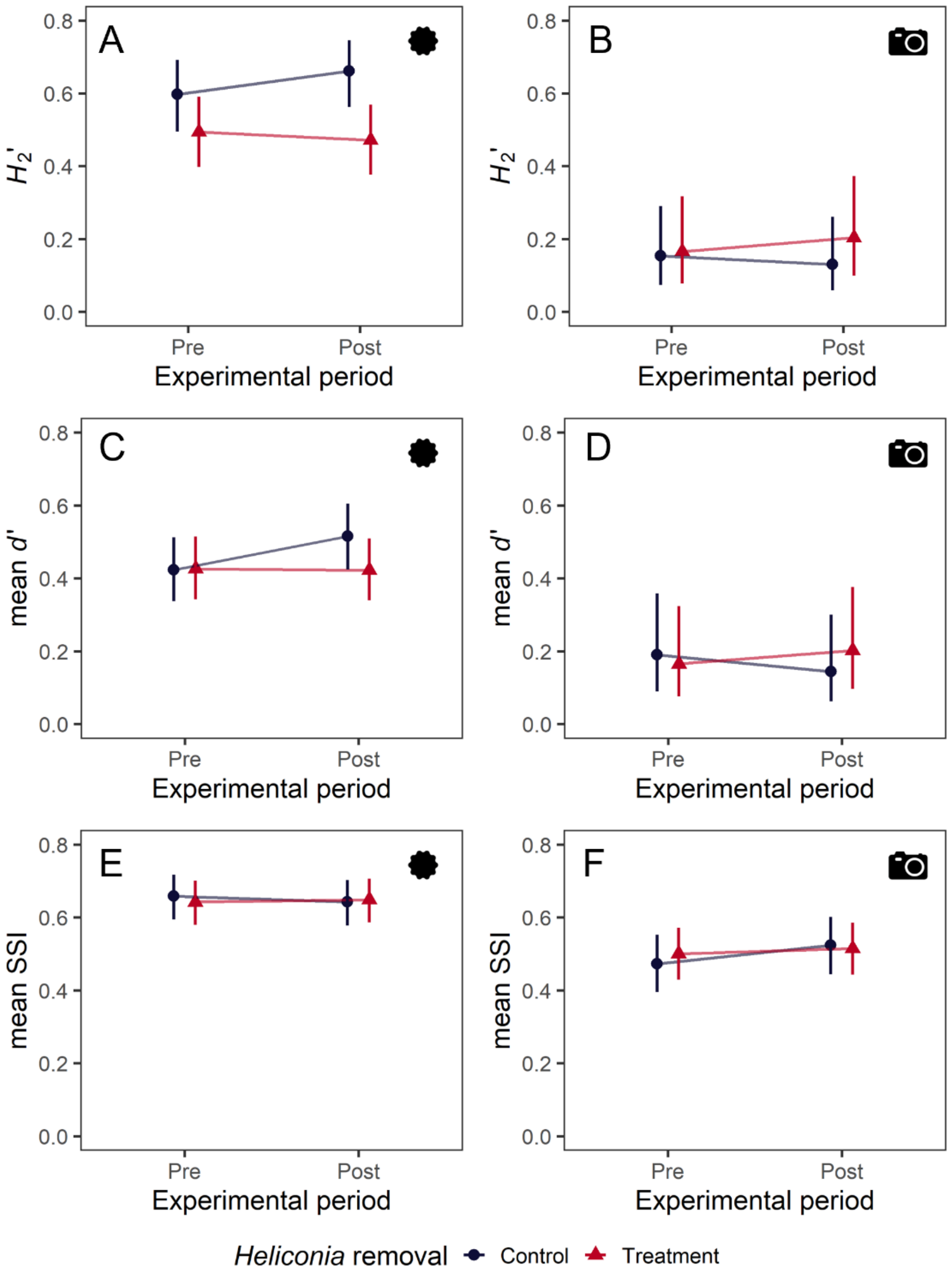


**Figure S4.** Effects of experimental Heliconia removal on species- and network-level specialization, measured as niche partitioning ($H_2$', *d*') and niche breadth (Species Specificity Index). Results are shown for pollen networks (left column) and camera networks (right column). Indices of species specialization were calculated from the hummingbird perspective and then averaged across all hummingbird species. Estimated means from GLMMs are presented alongside 95% confidence intervals; conceptually, a treatment effect is indicated by non-parallel lines.

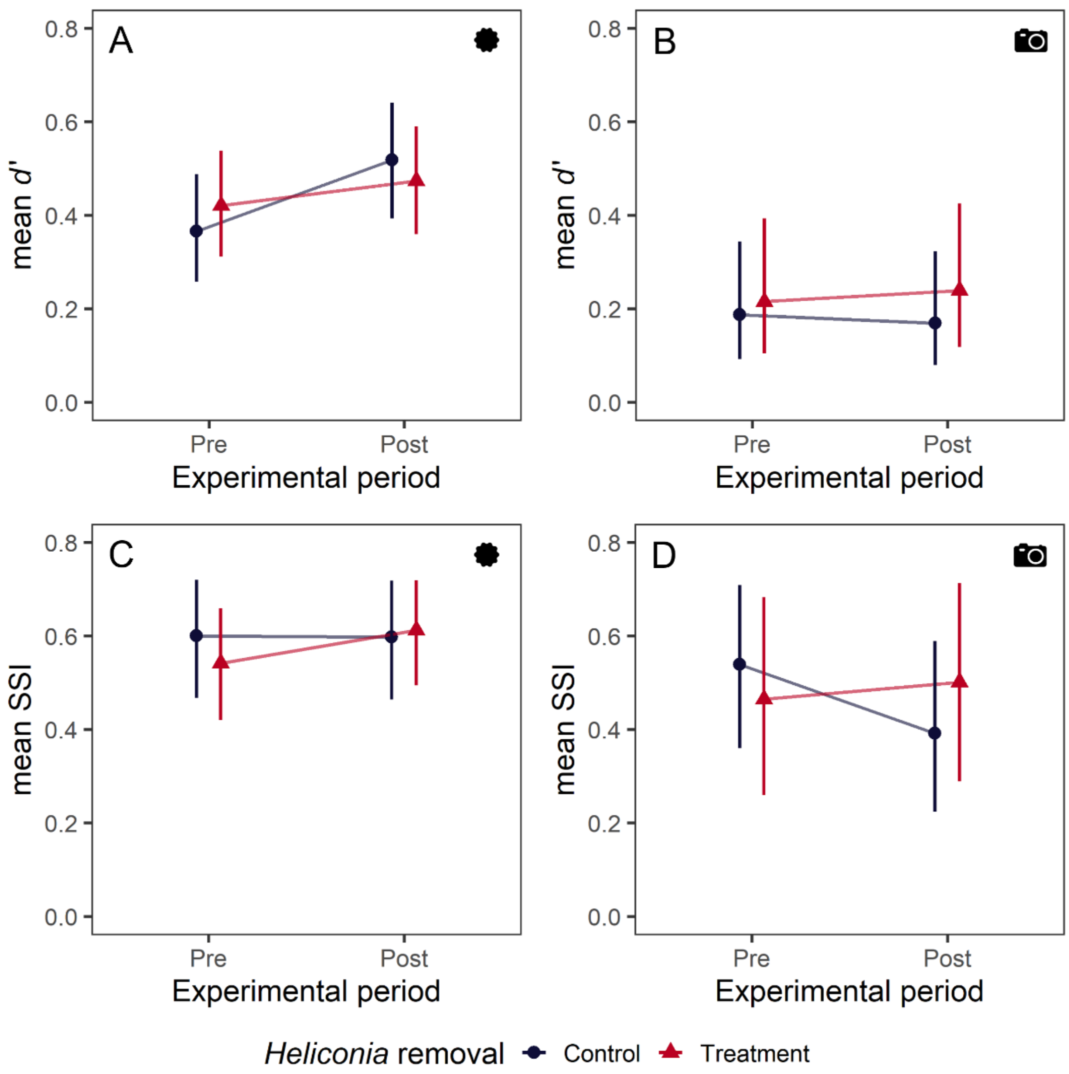


**Figure S5.** Effects of experimental *Heliconia* removal on species-level specialization, measured as niche partitioning (*d*') and niche breadth (Species Specificity Index). Results are shown for pollen networks (left column) and camera networks (right column). Indices of species specialization were calculated from the hummingbird perspective and then averaged across a *Heliconia* specialists (green hermits and violet sabrewings only). Estimated means from GLMMs are presented alongside 95% confidence intervals; conceptually, a treatment effect is indicated by non-parallel lines.

**Table S1.** Sample size per analysis. For all metrics except interaction turnover, values were calculated for both experimental periods (pre and post). For example, the analysis of individual specialization (recaptures) included 27 individuals each sampled twice, yielding 54 total observations.

| | | | All species | | | *Heliconia* specialists | | |
|---|---|---|---|---|---|---|---|---|
| | Level | Metric | Total rows | # control replicates | # treatment replicates | Total rows | # control replicates | # treatment replicates |
| Pollen | Network | $H_2$’ | 30 | 8 | 7 | | | |
| | Species | mean *d*’ | 32 | 8 | 8 | 30 | 8 | 7 |
| | Species | mean SSI | 34 | 8 | 9 | 30 | 8 | 7 |
| | Individual (all) | # morphotypes/bird | 302 | 11 | 13 | 126 | 10 | 13 |
| | Individual (recaptures) | # morphotypes/bird | 54 | 8 | 8 | 32 | 6 | 5 |
| Camera | Network | interaction turnover | 30 | 15 | 15 | | | |
| | Network | $H_2$’ | 60 | 14 | 16 | | | |
| | Species | mean *d*’ | 60 | 14 | 16 | 58 | 13 | 16 |
| | Species | mean SSI | 60 | 14 | 16 | 58 | 13 | 16 |

**Table S2.** Number of hours of video footage and pollen samples included in each dataset.

| Dataset | Purpose | Camera | | Pollen | |
|---|---|---|---|---|---|
| | | Data excluded | Number of hours | Data excluded | Number of samples |
| Full dataset | Divide into subsets for purposes detailed below | Dates without any flowers visible on camera[1] | 20,735 | None | 307 |
| Normal visitation | Understand 'normal' visitation patterns in this study system<br><br>Create pollen and camera meta-networks | Videos from 'post' period of treatment replicates | 15,119 | Samples from 'post' period of treatment replicates | 228 |
| Experimental | Examine hummingbird responses to experimental *Heliconia* removal | Cameras without videos for both experimental periods (pre and post) | 19,870 | Replicates without samples for both experimental periods (pre and post) | 302 |
| Sampling method | Examine how sampling method (pollen *vs.* camera) influences specialization<br><br>Examine the correlation between networks sampled with different methods | Cameras without videos for both experimental periods (pre and post)<br><br>Videos from 'post' period | 9,603 | Samples from 'post' period | 163 |

1, Some plant species did not produce open flowers every day

**Table S3**. Contrasts from GLMMs examining how experimental *Heliconia* removal (treatment) influences ecological specialization at the network, species, and individual level. Contrasts were calculated using emmeans (Lenth, 2020) and can be interpreted on the data scale; for instance, compared to control replicates, treatment replicates had 1.45 times more pre-to-post change in pollen network specialization measured as $H_2$’ (95%: 0.58-3.63).

| | | | All species | | | *Heliconia* specialists | | |
|---|---|---|---|---|---|---|---|---|
| | Specialization level | Specialization metric | Ratio[1] | 2.5% | 97.5% | Ratio[1] | 2.5% | 97.5% |
| Pollen | Network | | 1.45 | 0.58 | 3.63 | | | |
| | Species | mean *d*’ | 1.61 | 0.65 | 3.99 | 1.24 | 0.43 | 3.54 |
| | Species | mean SSI | 0.93 | 0.72 | 1.2 | 1.49 | 0.72 | 3.07 |
| | Individual: all individuals | # morphotypes/bird | 1.15 | 0.87 | 1.51 | 1.17 | 0.79 | 1.73 |
| | Individual: recaptures only | # morphotypes/bird | 1.78 | 1.04 | 3.06 | 1.74 | 1.16 | 2.6 |
| Camera | Network | $H_2$’ | 0.89 | 0.68 | 1.18 | | | |
| | Species | mean *d*’ | 0.82 | 0.65 | 1.04 | 0.81 | 0.53 | 1.23 |
| | Species | mean SSI | 1.05 | 0.94 | 1.18 | 1.17 | 0.88 | 1.55 |

**Statistical model**: specialization metric ~ treatment * experimental period + (1|site/replicate)

1, (Post:Pre in Treatment) / (Post:Pre in Control)

**Table S4.** Summary of GLMMs examining how experimental *Heliconia* removal (treatment) influences the number of pollen morphotypes per captured hummingbird. 'Recaptures' refers to only the hummingbirds captured during both capture sessions (pre and post); these models included an additional random effect for Bird ID. Wald confidence intervals, Wald *P*-values, and random effect standard deviations are provided. All models used a truncated generalized Poisson (log link), and values presented are on scale of the link function.

| | **POLLEN PER INDIVIDUAL** | All species | | | | | *Heliconia* specialists | | | | |
|---|---|---|---|---|---|---|---|---|---|---|---|
| | | Est | 2.5% | 97.5% | *z* | *P* | Est | 2.5% | 97.5% | *z* | *P* |
| All individuals | (Intercept) | 0.87 | 0.64 | 1.1 | 7.35 | <0.001 | 0.86 | 0.56 | 1.17 | 5.6 | <0.001 |
| | Treatment | 0.03 | -0.27 | 0.33 | 0.21 | 0.83 | 0.07 | -0.31 | 0.44 | 0.34 | 0.735 |
| | Post | -0.17 | -0.38 | 0.04 | -1.57 | 0.117 | -0.17 | -0.45 | 0.11 | -1.18 | 0.238 |
| | Treatment x Post | 0.14 | -0.14 | 0.41 | 0.99 | 0.32 | 0.16 | -0.23 | 0.54 | 0.8 | 0.425 |
| | (Random Intercept) Site | 0.00 | | | | | 0.00 | | | | |
| | (Random Intercept) Replicate | 0.27 | | | | | 0.32 | | | | |
| Recaptures only ($N_{All}$ = 27, $N_{GV}$ = 16) | (Intercept) | 0.93 | 0.67 | 1.2 | 6.97 | <0.001 | 1.09 | 0.86 | 1.31 | 9.63 | <0.001 |
| | Treatment | -0.15 | -0.5 | 0.21 | -0.8 | 0.426 | -0.25 | -0.54 | 0.05 | -1.63 | 0.104 |
| | Post | -0.34 | -0.73 | 0.05 | -1.69 | 0.092 | -0.35 | -0.63 | -0.08 | -2.5 | 0.013 |
| | Treatment x Post | 0.58 | 0.05 | 1.1 | 2.16 | 0.031 | 0.55 | 0.17 | 0.94 | 2.84 | 0.004 |
| | (Random Intercept) Site | 0.09 | | | | | 0.18 | | | | |
| | (Random Intercept) Replicate | 0.00 | | | | | 0.00 | | | | |
| | (Random Intercept) Bird | 0.00 | | | | | 0.00 | | | | |

**Table S5.** Summary of GLMMs examining how experimental *Heliconia* removal (treatment) influences three metrics of ecological specialization, calculated from pollen networks for each experimental period (pre and post). Wald confidence intervals, Wald *P*-values, and random effect standard deviations are provided. All models used a beta distribution (logit link), and values presented are on the scale of the link function.

| | **POLLEN NETWORKS** | All species | | | | | *Heliconia* specialists | | | | |
|---|---|---|---|---|---|---|---|---|---|---|---|
| | | Est | 2.5% | 97.5% | *z* | *P* | Est | 2.5% | 97.5% | *z* | *P* |
| $H_2$’ | (Intercept) | -1.71 | -2.48 | -0.94 | -4.34 | <0.001 | | | | | |
| | Treatment | 0.09 | -1.01 | 1.19 | 0.16 | 0.872 | | | | | |
| | Post | -0.19 | -0.93 | 0.55 | -0.5 | 0.618 | | | | | |
| | Treatment x Post | 0.44 | -0.59 | 1.48 | 0.84 | 0.401 | | | | | |
| | (Random Intercept) Site | 0.00 | | | | | | | | | |
| | (Random Intercept) Replicate | 0.81 | | | | | | | | | |
| mean *d*’ | (Intercept) | -1.45 | -2.28 | -0.63 | -3.44 | 0.001 | -1.47 | -2.24 | -0.69 | -3.7 | <0.001 |
| | Treatment | -0.17 | -1.33 | 0.99 | -0.29 | 0.775 | 0.17 | -0.81 | 1.16 | 0.34 | 0.732 |
| | Post | -0.33 | -1.09 | 0.43 | -0.85 | 0.393 | -0.13 | -1 | 0.74 | -0.29 | 0.772 |
| | Treatment x Post | 0.58 | -0.46 | 1.62 | 1.09 | 0.275 | 0.27 | -0.99 | 1.52 | 0.42 | 0.677 |
| | (Random Intercept) Site | 0.00 | | | | | 0.65 | | | | |
| | (Random Intercept) Replicate | 0.93 | | | | | 0.00 | | | | |
| mean SSI | (Intercept) | -0.11 | -0.41 | 0.2 | -0.7 | 0.485 | 0.16 | -0.54 | 0.85 | 0.44 | 0.657 |
| | Treatment | 0.11 | -0.28 | 0.5 | 0.56 | 0.575 | -0.3 | -1.43 | 0.84 | -0.52 | 0.606 |
| | Post | 0.2 | -0.15 | 0.56 | 1.12 | 0.265 | -0.6 | -1.46 | 0.27 | -1.35 | 0.176 |
| | Treatment x Post | -0.15 | -0.63 | 0.34 | -0.59 | 0.558 | 0.74 | -0.55 | 2.04 | 1.13 | 0.259 |
| | (Random Intercept) Site | 0.13 | | | | | 0.35 | | | | |
| | (Random Intercept) Replicate | 0.15 | | | | | 0.33 | | | | |

**Table S6.** Summary of GLMMs examining how experimental *Heliconia* removal (treatment) influences three metrics of ecological specialization, calculated from camera networks for each experimental period (pre and post). Wald confidence intervals, Wald *P*-values, and random effect standard deviations are provided. All models used a beta distribution (logit link), and values presented are on the scale of the link function.

| | **CAMERA NETWORKS** | All species | | | | | *Heliconia* specialists | | | | |
|---|---|---|---|---|---|---|---|---|---|---|---|
| | | Est | 2.5% | 97.5% | *z* | *P* | Est | 2.5% | 97.5% | *z* | *P* |
| $H_2'$ | (Intercept) | 0.39 | -0.01 | 0.79 | 1.89 | 0.059 | | | | | |
| | Treatment | -0.46 | -1 | 0.08 | -1.69 | 0.092 | | | | | |
| | Post | 0.28 | -0.17 | 0.73 | 1.21 | 0.228 | | | | | |
| | Treatment x Post | -0.29 | -0.9 | 0.32 | -0.94 | 0.347 | | | | | |
| | (Random Intercept) Site | 0.10 | | | | | | | | | |
| | (Random Intercept) Replicate | 0.45 | | | | | | | | | |
| mean *d'* | (Intercept) | -0.31 | -0.66 | 0.04 | -1.74 | 0.082 | -0.57 | -1.06 | -0.08 | -2.26 | 0.02 |
| | Treatment | 0.01 | -0.41 | 0.44 | 0.07 | 0.948 | 0.21 | -0.32 | 0.74 | 0.77 | 0.44 |
| | Post | 0.37 | 0.08 | 0.67 | 2.46 | 0.014 | 0.63 | 0.09 | 1.16 | 2.29 | 0.02 |
| | Treatment x Post | -0.37 | -0.78 | 0.04 | -1.76 | 0.078 | -0.37 | -1.09 | 0.35 | -1.01 | 0.31 |
| | (Random Intercept) Site | 0.32 | | | | | 0.56 | | | | |
| | (Random Intercept) Replicate | 0.41 | | | | | 0.00 | | | | |
| mean SSI | (Intercept) | 0.68 | 0.42 | 0.94 | 5.1 | <0.001 | 0.42 | -0.09 | 0.93 | 1.63 | 0.104 |
| | Treatment | -0.1 | -0.41 | 0.21 | -0.6 | 0.545 | -0.27 | -0.86 | 0.32 | -0.9 | 0.367 |
| | Post | -0.07 | -0.31 | 0.17 | -0.57 | 0.566 | -0.01 | -0.51 | 0.49 | -0.04 | 0.971 |
| | Treatment x Post | 0.15 | -0.18 | 0.47 | 0.88 | 0.376 | 0.37 | -0.3 | 1.04 | 1.08 | 0.279 |
| | (Random Intercept) Site | 0.25 | | | | | 0.46 | | | | |
| | (Random Intercept) Replicate | 0.26 | | | | | 0.41 | | | | |

**Table S7.** Sampling completeness summarized by sampling method, across all networks (pre and post). Sample size *N* refers to the number of networks included. Within the camera networks, two replicates had particularly large pre-to-post differences in sampling completeness and were considered outliers.

| Pollen | | | Camera | | | Camera (without outliers) | | |
|---|---|---|---|---|---|---|---|---|
| Mean | SD | *N* | Mean | SD | *N* | Mean | SD | *N* |
| 0.5 | 0.19 | 49 | 0.91 | 0.11 | 64 | 0.92 | 0.08 | 60 |

**Table S8.** Contrasts from GLMMs examining whether sampling completeness differs between experimental periods and/or with *Heliconia* removal treatment. For pollen networks, a pre-to-post change in sampling completeness was detected in treatment replicates, indicated by 95% confidence intervals that did not overlap zero (bolded values in table). These networks were not included in the analysis of interaction turnover.

| | Pollen | | | Camera | | | Camera (without outliers) | | |
|---|---|---|---|---|---|---|---|---|---|
| | Ratio[1] | 2.5% | 97.5% | Ratio[1] | 2.5% | 97.5% | Ratio[1] | 2.5% | 97.5% |
| Control | 0.88 | 0.66 | 1.18 | 1 | 0.95 | 1.06 | 0.98 | 0.94 | 1.03 |
| Treatment | **0.72** | **0.53** | **0.99** | 1.05 | 0.99 | 1.12 | 1.03 | 0.98 | 1.09 |

**Statistical model**: completeness ~ treatment * experimental period + (1|site/replicate)

1, Post:Pre

**Table S9.** Summary of GLMMs examining how experimental *Heliconia* removal (treatment) influences pre-to-post interaction turnover in camera networks. Wald confidence intervals, Wald *P*-values, and random effect standard deviations are provided. All models used a beta distribution (logit link), and values presented are on the scale of the link function.

| | | Binary (Whittaker's beta diversity) | | | | | Quantitative (Bray-Curtis dissimilarity) | | | | |
|---|---|---|---|---|---|---|---|---|---|---|---|
| | | Est | 2.5 % | 97.5% | *z* | *P* | Est | 2.5% | 97.5% | *z* | *P* |
| Total interaction turnover | (Intercept) - Control | -0.85 | -1.17 | -0.53 | -5.16 | <0.001 | -0.66 | -0.91 | -0.41 | -5.14 | <0.001 |
| | Treatment | -0.25 | -0.72 | 0.21 | -1.08 | 0.281 | 0.27 | -0.08 | 0.62 | 1.5 | 0.133 |
| | (Random Intercept) Site | 0.00 | | | | | 0.00 | | | | |
| Species turnover | (Intercept) - Control | -1.52 | -2 | -1.04 | -6.19 | <0.001 | -2.23 | -2.68 | -1.78 | -9.74 | <0.001 |
| | Treatment | -0.35 | -1 | 0.3 | -1.05 | 0.293 | -0.31 | -0.91 | 0.29 | -1.01 | 0.311 |
| | (Random Intercept) Site | 0.00 | | | | | 0.00 | | | | |
| Turnover among shared species | (Intercept) - Control | -2.09 | -2.44 | -1.73 | -11.46 | <0.001 | -1.08 | -1.28 | -0.88 | -10.56 | <0.001 |
| | Treatment | 0.47 | 0.03 | 0.9 | 2.09 | 0.037 | 0.5 | 0.23 | 0.77 | 3.64 | <0.001 |
| | (Random Intercept) Site | 0.24 | | | | | 0.00 | | | | |

**Table S10.** Contrasts from GLMMs examining how experimental *Heliconia* removal influences pre-to-post interaction turnover in camera networks. Contrasts were calculated using emmeans (Lenth, 2020) and can be interpreted on the data scale; for instance, compared to control replicates, treatment replicates had 1.49 times more interaction turnover among species present in both experimental periods (95% CI: 1.01-2.22).

| | Binary (Whittaker's beta diversity) | | | Quantitative (Bray-Curtis dissimilarity) | | |
|---|---|---|---|---|---|---|
| | Ratio[1] | 2.5% | 97.5% | Ratio[1] | 2.5% | 97.5% |
| Total interaction turnover | 0.83 | 0.59 | 1.18 | 1.18 | 0.94 | 1.49 |
| Species turnover | 0.74 | 0.42 | 1.33 | 0.75 | 0.42 | 1.34 |
| Turnover among shared species | 1.49 | 1.01 | 2.22 | 1.42 | 1.16 | 1.73 |

**Statistical model:** interaction turnover ~ treatment + (1|site)

1, Treatment:Control

**Table S11.** Contrasts from GLMMs examining how sampling method (pollen *versus* camera) influences three metrics of ecological specialization calculated from 40 networks sampling during parallel time periods (20 pollen networks, 20 camera networks). Contrasts were calculated using emmeans (Lenth, 2020) and can be interpreted on the data scale; for instance, when specialization is measured as $H_2$', camera networks are ~3 times more specialized than pollen networks (95%: 2.08-4.48 times).

| | Ratio[1] | 2.5% | 97.5% |
|---|---|---|---|
| $H_2$' | 3.05 | 2.08 | 4.48 |
| mean *d*' | 2.02 | 1.33 | 3.05 |
| mean SSI | 1.43 | 1.22 | 1.69 |
| **Statistical model:** specialization metric ~ sampling method + (1\|site/replicate) | | | |

1, Camera:Pollen

**Table S12.** Summary of GLMMs examining how sampling method (pollen *versus* camera) influences three metrics of ecological specialization calculated from 40 networks sampling during parallel time periods (20 pollen networks, 20 camera networks). Wald confidence intervals, Wald *P*-values, and random effect standard deviations are provided. All models used a beta distribution (logit link), and values presented are on the scale of the link function.

| | | Est | 2.5% | 97.5% | *z* | *P* |
|---|---|---|---|---|---|---|
| $H_2$' | (Intercept) – Pollen network | -1.6 | -2.04 | -1.15 | -7.03 | <0.001 |
| | Camera network | 1.65 | 1.15 | 2.16 | 6.38 | <0.001 |
| | (Random Intercept) Site | 0.33 | | | | |
| | (Random Intercept) Replicate | 0.00 | | | | |
| mean *d*' | (Intercept) – Pollen network | -1.25 | -1.74 | -0.77 | -5.07 | <0.001 |
| | Camera network | 1.04 | 0.5 | 1.59 | 3.77 | <0.001 |
| | (Random Intercept) Site | 0.00 | | | | |
| | (Random Intercept) Replicate | 0.00 | | | | |
| mean SSI | (Intercept) – Pollen network | -0.19 | -0.45 | 0.06 | -1.48 | 0.139 |
| | Camera network | 0.8 | 0.46 | 1.15 | 4.6 | <0.001 |
| | (Random Intercept) Site | 0.00 | | | | |
| | (Random Intercept) Replicate | 0.22 | | | | |

**Table S13.** Summary of GLMMs examining the relationship between pollen and camera networks collected at the same site during the same time period. Wald confidence intervals, Wald *P*-values, and random effect standard deviations are provided. All models used a beta distribution (logit link), and values presented are on the scale of the link function.

| | | Est | 2.5% | 97.5% | *z* | *P* |
|---|---|---|---|---|---|---|
| $H_2$' | (Intercept) | -1.75 | -3.12 | -0.38 | -2.5 | 0.012 |
| | Camera network | 0.57 | -1.85 | 3 | 0.46 | 0.643 |
| | (Random Intercept) Site | 0.00 | | | | |
| | pseudo-$R^2$ | 0.014 | | | | |
| mean *d*' | (Intercept) | -0.92 | -2.34 | 0.49 | -1.28 | 0.202 |
| | Camera network | -0.74 | -3.72 | 2.24 | -0.48 | 0.628 |
| | (Random Intercept) Site | 0.00 | | | | |
| | pseudo-$R^2$ | 0.014 | | | | |
| mean SSI | (Intercept) | 0.67 | -0.77 | 2.12 | 0.91 | 0.361 |
| | Camera network | -1.08 | -3.27 | 1.11 | -0.97 | 0.332 |
| | (Random Intercept) Site | 0.00 | | | | |
| | pseudo-$R^2$ | 0.048 | | | | |
